# Short-term plasticity at Purkinje to deep cerebellar nuclear neuron synapses supports a slow gain-control mechanism enabling scaled linear encoding over second-long time windows

**DOI:** 10.1101/749259

**Authors:** Christine M. Pedroarena

**Affiliations:** Department for Cognitive Neurology, Hertie-Institute for Clinical Brain Research, 72076 Tübingen, Germany; Systems Neurophysiology, Werner Reichardt Center for Integrative Neuroscience, University Tübingen, 72076 Tübingen, Germany

## Abstract

Modifications in the sensitivity of neural elements allow the brain to adapt its functions to varying demands. Frequency-dependent short-term synaptic depression (STD) provides a dynamic gain-control mechanism enabling adaptation to different background conditions alongside enhanced sensitivity to input-driven changes in activity. In contrast, synapses displaying frequency-invariant transmission can faithfully transfer ongoing presynaptic rates enabling linear processing, deemed critical for many functions. However, rigid frequency-invariant transmission may lead to runaway dynamics and low sensitivity to changes in rate. Here, I investigated the Purkinje cell to deep cerebellar nuclei neuron synapses (PC_DCNs), which display frequency-invariance, and yet, PCs maintain background-activity at disparate rates, even at rest. Using protracted PC_DCNs activation (120s) in cerebellar slices to mimic background-activity, I identified a previously unrecognized frequency-dependent, slow STD (S_STD) of PC_DCN inhibitory postsynaptic currents. S_STD supports a novel form of gain-control that enabled—over second-long time windows—scaled linear encoding of PC rate changes mimicking behavior-driven/learned PC-signals, alongside adaptation to background-activity. Cell-attached DCN recordings confirmed these results. Experimental and computational modeling results suggest S_STD-gain-control may emerge through a slow depression factor combined with balanced fast-short-term plasticity. Finally, evidence from opto-genetic experiments, statistical analysis and computer simulations pointed to a presynaptic, input-specific and possibly activity-dependent decrease in active synaptic release-sites as the basis for S_STD. This study demonstrates a novel slow gain-control mechanism, which could explain efficient and comprehensive PC_DCN linear transfer of input-driven/learned PC rates over behavioral-relevant time windows despite disparate background-activity, and furthermore, provides an alternative pathway to hone PCs output via background-activity control.

**SIGNIFICANCE STATEMENT:** The brain can adapt to varying demands by dynamically changing the gain of its synapses; however, some tasks require linear transfer of presynaptic rates over extended periods, seemingly incompatible with non-linear gain adaptation. Here, I report a novel gain-adaptation mechanism, which enables scaled linear encoding of changes in presynaptic rates over second-long time windows and adaptation to background-activity at longer time-scales at the Purkinje to deep cerebellar nuclear neurons synapses (PC_DCNs). A previously unrecognized PC_DCN slow and frequency-dependent short-term synaptic depression (S_STD), together with frequency-invariant transmission at faster time scales likely explains this process. This slow-gain-control/modulation mechanism may enable efficient linear encoding of second-long presynaptic signals under diverse synaptic background-activity conditions, and flexible fine-tuning of synaptic gains by background-activity modulation.

## INTRODUCTION

Changes in the sensitivity or gain of neural structures is a general principle that allows the brain to adapt to varying sensory, motor, emotional, or cognitive demands ^1,2^. One mechanism supporting gain adaptation is use-dependent short-term synaptic depression (STD), where high levels of activity lead to lower synaptic strength (idem), thereby dynamically adapting the output while enhancing the sensitivity to changes in presynaptic activity ^3,4^. The latter is because the change in synaptic strength lags behind the change in presynaptic firing rate, such that for a short time window —in the millisecond range for fast-STDs ^5^ — the output becomes proportional to the new rate, before readapting. This process produces brief transient signals informative about the timing and proportion of the initial change in rate ^3^. However, beyond this brief window, STD filters the synaptic output preventing faithful transfer of complete presynaptic rate profiles, which may hold important information.

For instance, the cerebellar cortex principal neurons, the GABAergic Purkinje cells (PCs), display changes in rate associated to cerebellar-controlled behaviors(e.g. eye-blinks, saccades, and whisks) extending beyond the referred STD time-window, which are considered critical for the representation and control of specific behavioral parameters (e.g. ^6–9^). Suitably, the synapses from the PCs to their main target, the deep-cerebellar-nuclei-neurons (PC_DCNs) in fact, display frequency-invariant transmission supporting linear encoding of PC rates ^10^. Mechanistically, PC_DCN frequency-invariance is likely mediated by a developmentally-regulated synaptotagmin7-dependent synaptic facilitation ^11^ counteracting a frequency-dependent fast-STD ^12^. Functionally, linear encoding has been widely reported in the cerebellum, including the cerebellar nuclei ^6,7, 13–17^, but see ^18^. Reasonably, the latter studies focused on time-windows matching usual behaviors, i.e. ranging from sub-second to few seconds.

Yet, PCs discharge simple spikes almost continuously at disparate rates diverging by as much as hundred Hz, even in awake animals at rest and within the same region ^19–25^. Therefore, PC signals associated with particular behaviors consist in transitory changes in rate embedded in the background-activity. PC background-activity has received much less attention than the behavior-driven changes in PC rate, however, it is known it might be altered in cerebellar associated diseases ^26,27^ and that diversity and variability in PC background rates originate in differences in intrinsic properties and ongoing PC inputs ^14,23,28^. This suggests that PC background activity might be a controlled variable whose integrity might be required for correct cerebellar function, potentially moldable via PC intrinsic plasticity ^29,30^, plastic changes of the upstream cerebellar circuits ^31^, and/or context-dependent changes in the ongoing activity of PCs numerous inputs. However, PC background-activity presents a challenge for the reliable transfer of PC behavior-related signals using frequency-invariant synapses because the evoked changes in firing rate can be masked by the large span of PCs rates converging on DCNs ^3^; and for keeping DCNs within their working-firing range.

Coexpression of different forms of short-term plasticity (STP) expands the computational power of single synapses ^10,32–34^; however, PC_DCN-STP has not been previously explored at timescales matching background-activity. This study, by closing this gap, aimed to identify STP forms and mechanisms conciliatory of background-activity processing with linear encoding over behaviour-relevant time-windows.

## RESULTS

### Slow, frequency dependent short-term depression at PC_DCNs

To explore the susceptibility of PC_DCNs to changes in sustained activity, here I recorded large, putative projecting glutamatergic DCNs ^35^ in cerebellar slices from juvenile (P21-P32) or adult (P55_P72) mice under conditions alike those found *in situ* (i.e. temperature: 36.5°C ± 0.5°C and extracellular CaCl_2_ concentration: 1.5 mM) using whole cell patch clamp or cell attached modes. To mimic the persistent discharge of PCs, I used protracted electrical stimulation of PC axons in the surrounding white matter (≥ 2 minutes) at three different stimulating background frequencies (BF: 10, 29.5 and 67 Hz), which are within the PCs firing rate range in awake animals at rest ^19,23^.

The responses to 2 minutes-long trains of stimuli confirmed an initial phase of rapid decrease in phasic IPSCs peak amplitude with successive events ^10,12,36^ which tends to slow down and level after tens of events ^10^ (Figure 1A-B). However, a different slow phase of decay in IPSC amplitude ensuing the fast one was unveiled, most evident by analyzing the results in a compressed timescale (Figure 1B, insets show fits using double exponential decay functions).

**Figure 1.**
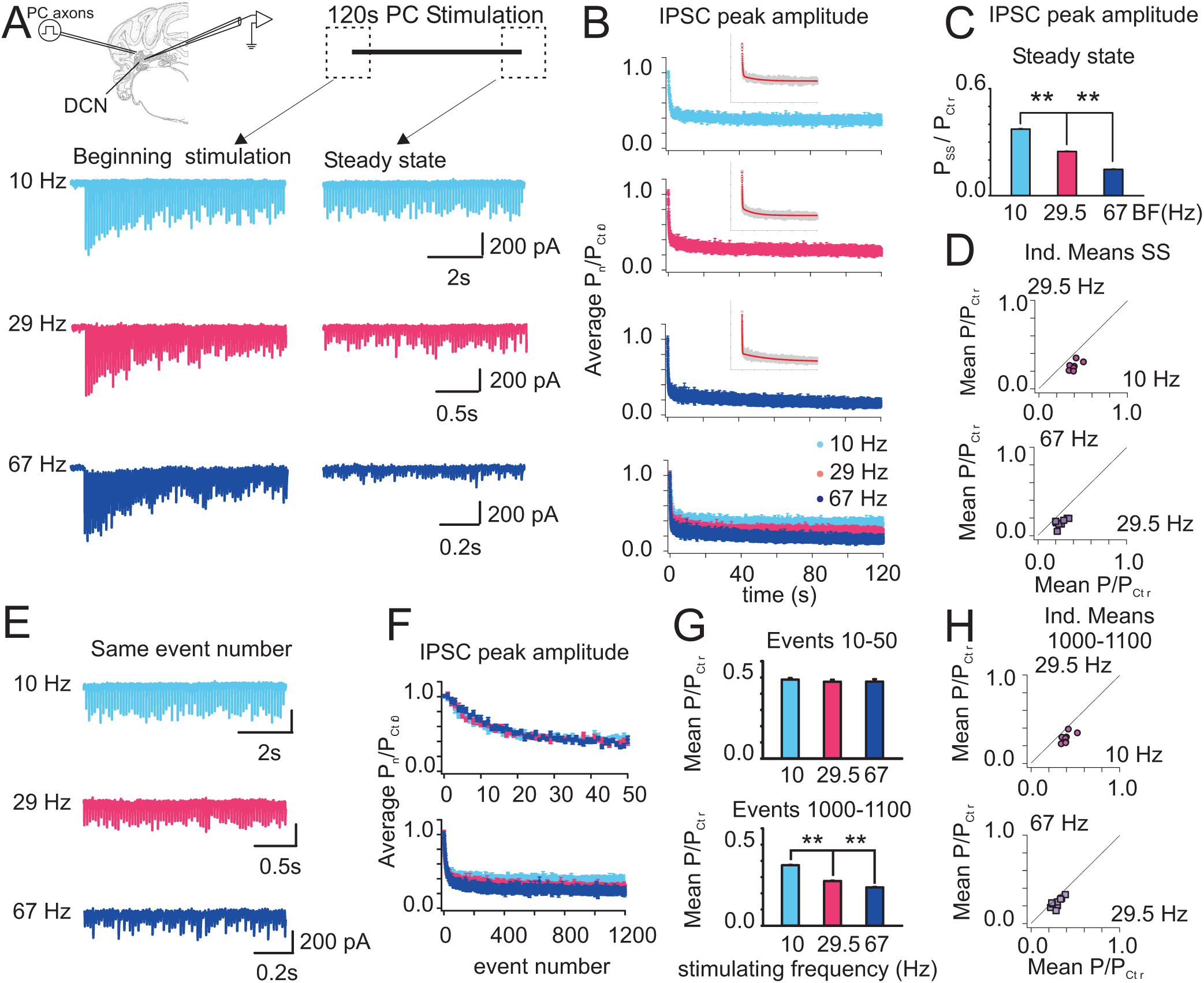
Protracted PC stimulation unveils a slow and frequency-dependent short-term depression. (A) Top: Sketch of experimental configuration and stimulation paradigm. Below: typical examples of IPSCs from the same neuron evoked by 2 minutes trains of stimuli applied to PC axons at the frequencies (BF) indicated on the left, at the beginning (right panel) and at the end of stimulation (left panel). Note the different time scales. (B) Top to bottom: Mean normalized IPSC peak amplitude as a function of time for BFs 10 Hz (light blue, n= 11), 29.5 Hz (red, n=9) and 67 Hz (dark blue, n=9), and all three plots overlaid (symbols represent mean ± SEM) illustrate two phases of decay. The double exponential decay fits in the insets: fast tau: 0.81±0.04s, 0.41±0.012s, 0.33±0.007s and slow tau: 23±1.9s, 18±0.5s, and 47±1.2s for 10, 29,5 and 67 Hz BFs respectively). (C) Average normalized steady-state phasic IPSCs amplitude (IPSC_SS_) vs. BF (averages calculated over the last one hundred events of the data plotted in Fig. 1B; 0.37 ± 0.002, 0.25 ± 0.002 and 0.15 ±0.002 for 10, 29.5 and 67 Hz BFs respectively, t-test:10 vs. 29.5 Hz and 29.5 versus 67Hz P<0.001 for both comparisons). (D) Results from individual neurons illustrate decreasing IPSC_SS_ with increasing BF: Top: IPSC amplitude for 29.5 Hz vs. 10 Hz; Bottom: idem but for 67 Hz vs. 29.5 Hz (details in main text). Note that all points fall below the unity line. (E) Typical examples from the same neuron in (A) of IPSCs evoked by the same ordinal stimuli but at different BF (indicated on the left). Note, the 10 Hz trace is the same shown in (A) for comparison. (F) IPSC amplitude as in (B) but as a function of the event number (color labeled as in (B)) Top: Events 1 to 50. Bottom: Events 1 to 1200. (G) Averages data shown in Fig. 1F calculated over the events indicated on top of each bar plot. Events 10 to 50: 0.49±0.01, 0.47±0.02 and 0.47±0.03, for 10, 29.5 and 67 Hz respectively; MWRS test: 10 vs. 29.5 Hz, P = 0.115; 29.5 vs. 67 Hz P= 0.7. E events 1000 to 1100: 0.37±0.002, 0.25± 0.002, 0.147± 0.002, for 10, 29.5 and 67 Hz respectively, t-test, 10 vs. 29.5 and 29.5 vs. 67 Hz, P< 0.001 for both. (H) Similar as in (D) but for mean IPSCs amplitudes calculated over stimuli 1000th to 1100th for each neuron: Top: 29.5Hz vs. 10 Hz; Bottom: 67 vs. 29.5 Hz. All data as mean ± SEM. (**) and (*) indicate P< 0.001 and P<0.05 respectively, for these and following plots.

Remarkably, the average synaptic strength at the end of the 2-minutes stimulation periods decreased with increasing frequency (Figure. 1A, 1C; e.g., 0.37 ± 0.002 vs. 0.15 ±0.002 for 10 and 67 Hz BFs respectively). Importantly, the population results reflect the results from single experiments, depicted individually in Figure 1D. Note that although the magnitude of depression varied amongst neurons, for each case the normalized steady state synaptic strength decreased with increasing BF. (Mean values: 0.4 ± 0.02 (10 Hz), vs. 0.26± 0.02 (29.5 Hz), n=8 paired t-test, P<0.001; 0.26±0.02 (29.5 Hz) vs. 0.15±0.015 (67 Hz), n=9, paired t-test P< 0.001). Therefore, protracted activation of PC_DCN synapses revealed a frequency-dependent slow phase of short-term depression (hereafter S_STD). Although the differences in synaptic strength for different BFs seem small normalized to control, note that the strength at 10 Hz was more than double that found using 67 Hz, suggesting functional relevance.

To explore the buildup of frequency-dependence along the stimulating train I analyzed the changes in synaptic strength as a function of the event number (Figure 1E-H). In agreement with previous results ^10^, the IPSCs amplitudes initially depressed significantly but on average with slightly lower rates with increasing BFs (Figure 1F top), while later, over the next 50-100 events, the IPSCs evoked with different BFs displayed similar amplitudes (Figs. 1F, top and 1G, top). However, after hundreds of events the IPSCs amplitudes for different BFs increasingly diverged (Figure 1F, bottom). Significant differences in average IPSC amplitudes as a function of the BFs could be detected considering events 1000th to 1100^th^ or further (Figure 1G, bottom). Moreover, the results from single experiments are in agreement with the population ones (Fig. 1H): in all but one case, the mean IPSCs amplitude decreased with increasing BF (0.402± 0.02 vs. 0.29± 0.02, n= 8, paired t-test, P<0.001 for 10 vs. 29.5Hz; and 0.29±0.02 vs. 0.24±0.02, for 29.5Hz vs. 67 Hz, n= 9 paired t-test, P=0.003).

Similar results arose using Poisson-like trains of stimuli with mean frequencies 10 or 70 Hz (Fig. 2A), ruling out that the usual alternation in awake animals of regular and irregular periods of firing ^16,37^ could result in recovery from depression during long intervals abolishing the frequency differences in synaptic strength found here.

**Figure 2.**
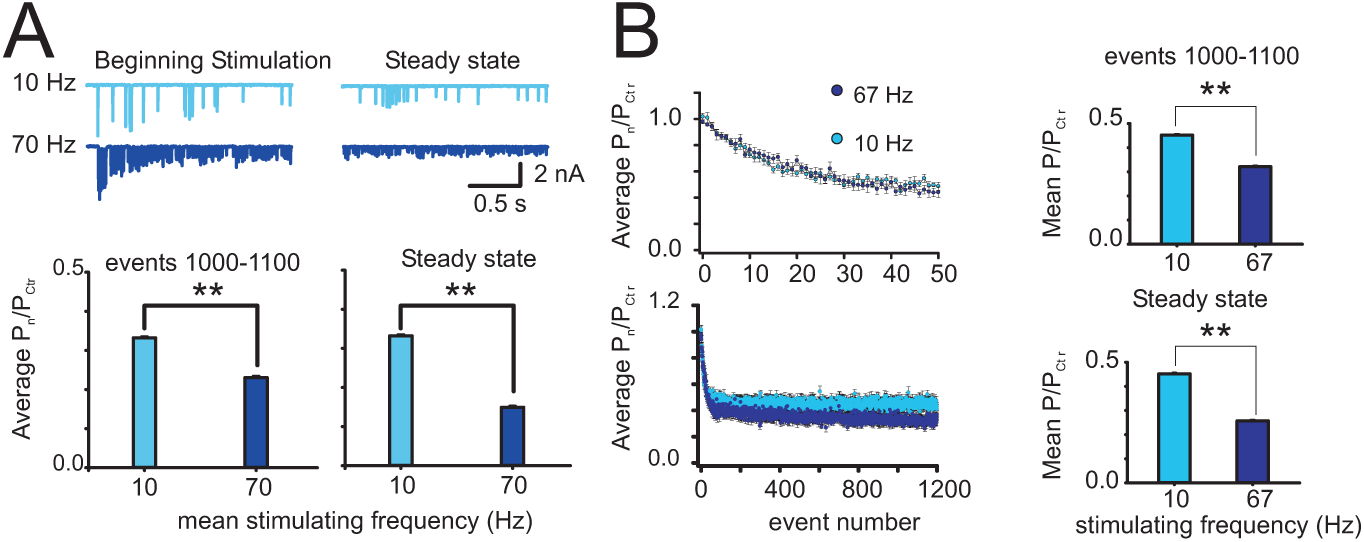
S_STD frequency dependence under irregular activation patterns and in adult mice. (A) Top traces: Representative traces of IPSCs evoked in the same neuron by 2 minutes Poisson-like stimulation at the mean frequencies indicated on the left. Bottom: Summary plots of mean IPSCs amplitudes calculated over the stimuli indicated on top of each graph as in Figure 1, but for Poisson like stimulation at 10 Hz (n=9) and 70 Hz (n=7). Significance assessed using Mann-Whitney rank sum test. (B) Left: Normalized average amplitude as a function of event number of PC-DCN IPSC recordings obtained from slices from P55 to P72 animals (BF 10 Hz, n=8, BF 67 Hz, n=7). Right: Summary plots of mean IPSC amplitudes calculated from the graphs on the left, over the events indicated on top of each bar-plot, (1000th to 1100th for 10 and 67 Hz BFs: 0.45±0.002 and 0.32±0.002 t-test P<0.001); (average of 100 events at steady state: 0.45±0.002 and 0.26±0.003 for 10 and 67 Hz respectively, t-test P<0.001).

Importantly, despite developmentally-regulated PC_DCN short-term plasticity ^10,11^, I found similar late phase of slow frequency-dependent decay in IPSCs amplitude using slices from adult mice (P55 to P72), albeit the levels of depression at steady state seem shallower than at younger ages (Fig. 2B). This strongly suggests that at young and adult PC_DCN synapses regular and irregular sustained activity induces frequency-dependent slow short-term depression (S_STD).

Due to the acknowledged difficulties in carrying DCN recordings from slices using older animals ^10,35^ and the need of particularly long recordings for the present study, the rest of the experiments were performed in slices from juvenile mice (> P20) as in other studies ^10,11^.

### Sustained PC activation modulates PC_DCN IPSCs evoked by changes in activation rate

Considering that PCs in awake-behaving animals typically display sustained activity, S_STD likely continuously controls the efficacy of PC terminals. This raises the question of whether and how S_STD modulates the responses to changes in PC rate associated to cerebellar controlled behaviors.

To address this question experimentally, PC_DCN synapses were first depressed to steady state using protracted stimulation at different BFs and afterwards, a brief, high frequency train of stimuli (HFS: 100 events, 180 Hz) was used to mimic behavior-driven transient signals. The HFS duration lays within the length of most common discrete motor events, e.g. saccades, reaches, steps, etc. To estimate and compare the synaptic output induced by HFSs preceded by different BFs, the total charge transferred during the HFS (Q_HFS_, *see Methods*), normalized to the charge transferred by corresponding control IPSCs and expressed per second was calculated. Analysis of changes in Q_HFS_ as a function of the preceding BF revealed a remarkable modulation (Fig. 3). Specifically, a decrease in Q_HFS_ with increasing BFs (Fig. 3A: no prior activity, 0Hz: 72±4.5, n= 22; 10 Hz: 61±3.3, n=16; 29.5Hz: 40±4.0, n=9; 67 Hz: 16±1.9, n=12; significance assessed using t-test: 0 vs. 10 Hz, P= 0.033, 1 tail; 10 vs. 29.5 Hz; P<0.001; 29.5 vs. 67 Hz, P<0.001). Note that the Q_HFS_ for HFSs preceded by 2 minutes stimulation at 67 Hz was approximately a quarter of the Q_HFS_ elicited by HFSs preceded by 10 Hz stimulation. These results support the view that the differences in steady state strength induced by different BFs have functional significance.

**Figure 3.**
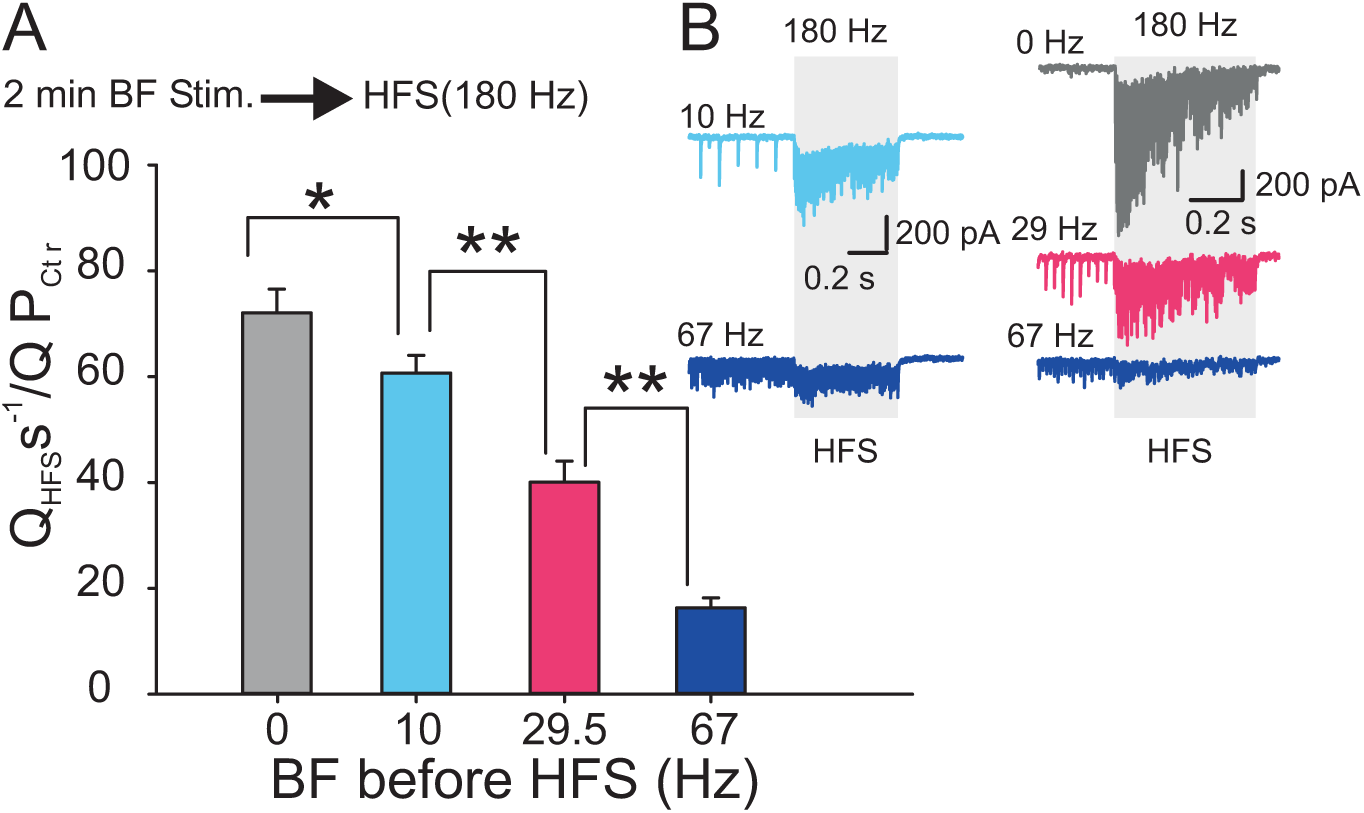
PC sustained activity modulates PC_DCN synaptic responses to PC high frequency stimulation trains (HFS) (A) Top: Stimulation paradigm. Bottom: Mean charge transfer amplitude during HFS (100 events, 180 Hz), normalized to that of control IPSCs as a function of prior background frequency. See main text for details. (B) Typical examples from two different neurons illustrating how prior BF (indicate on the left of each trace) modulates the amplitude of IPSCs evoked by the HFS (indicated by the grey box).

### Sustained PC activation modulates DCNs spiking responses evoked by changes in activation rate

Next, to verify the functional relevance of the latter finding, using cell attached recordings of spontaneously firing DCNs, I investigated the effect of similar stimulating protocols on DCNs spiking activity (Fig. 4). Under these conditions, the intracellular content was not dialyzed reflecting the conditions found in the intact animal/neuron. In addition, here, the same number of preceding stimuli was applied (7000 stimuli at 10 or 67 Hz), to rule out that the number of stimuli (and not the stimulating frequency) determined the changes in synaptic strength.

**Figure 4.**
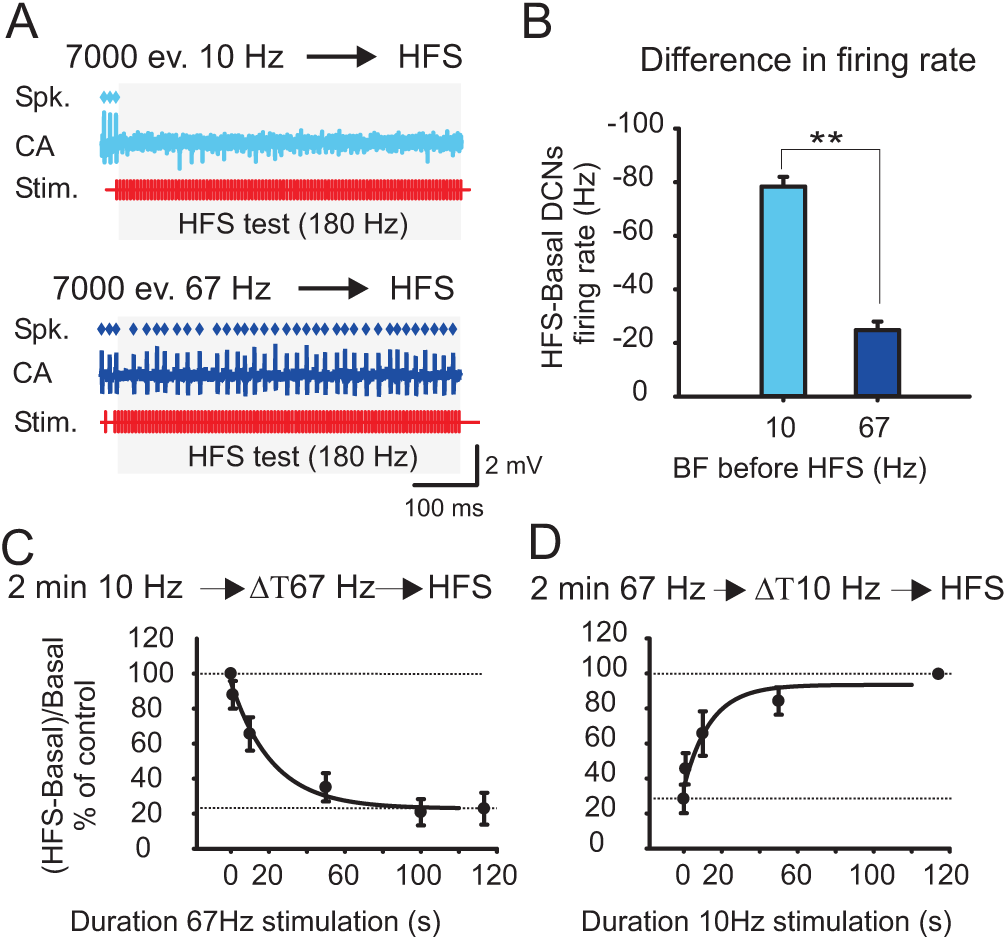
S_STD modulates HFSs suppression of DCN spontaneous firing rate (cell-attached recordings). (A) Top: example of response to a HFS delivered after 7000 stimuli at 10 Hz, Bottom idem after 7000 stimuli at 67 Hz (same unit). Spk.: time of spike occurrence; CA: cell-attached spike recording; Stim: time of stimulation. (B) Mean HFS-induced changes in DCNs firing rate (HFS-Basal rate) as a function of prior BF, n=6, P<0.001, t-test. (C) Effect of non-stationary PC stimulation on HFSs suppression of DCNs spontaneous activity: Top: stimulation paradigm; Bottom: Mean HFS-induced-changes in DCNs firing rate as a function of the 67 Hz train duration, normalized to responses evoked without switch to 67 Hz (upper dashed line, 100 %). Lower dashed line: the value of mean responses to HFS after 2min at 67 Hz only. The data was fitted with a single exponential function (R^2^: 0.99, n= 6). See text for details. (D) Idem, but for a switch in BF from 120s at 67 Hz to 10 Hz, (R^2^:0.94, n=6).

HFSs preceded by sustained stimulation at 10 Hz completely suppressed or sharply decreased DCNs firing rate, while after 67 Hz only a moderate effect was induced (Fig. 4A-B). These results strongly support the functional relevance of S_STD

Nest, since PCs ongoing firing rates vary with time, I asked how effective are non-stationary changes in PCs ongoing activity in modulating the responses to transient signals. Implicit to this question is the issue of the time course of S_STD. To address this question, using cell-attached recordings as in Fig. 4A, I interposed between the 2 minutes blocks of stimulation at 10 or 67 Hz and the HFS, a switch in the stimulating frequency (to 67 or 10 Hz respectively), for variable periods of time (shown 1, 10, 50 or 100s, Fig. 4C-D, see the scheme on top). The change in DCN firing rate for the 10-to-67-Hz-switches (normalized to the response to an HFS preceded by 120 s at 10 Hz only) decreased progressively with increasing periods of time under 67 Hz, with a time constant of 22 seconds. The change was undetectable for 67 Hz periods of 1 second or shorter and in average only after 100s, the suppression reached the same level than after 2 minutes of stimulation at 67 Hz. For the switch from 67 to 10 Hz, the time constant of recovery was 14 seconds, i.e., the recovery from deeper depression seemed faster than the buildup, but within a similar time range. Again, no influence could be detected for periods of 1 second or shorter (Fig. 4D).

The bi-directional amplitude modulation established in the course of tens of seconds highlight the time scale and flexibility associated to S_STD and consequent HFS response modulation. Importantly, the results revealed a time window of sub-second to few seconds, during which PC_DCN synaptic gain remains quasi stable. Remarkably, this is the range of durations of common motor events controlled by cerebellum.

### Preceding BF changes the gain but not the linear input/output transfer function of 0.5 s long signals

The above results suggested that for an interval of hundreds of milliseconds to seconds after a change in rate, the gain was independent of the new rate, possibly supporting linear encoding. For this to be the case, the mechanism of S_STD should be independent from the frequency invariance mechanism observed at shorter time scales. Alternatively, if S_STD results for instance from decreased facilitation or enhanced fast depression, the transfer function for changes in rate could show non-linearity. Therefore, to explore this issue I asked whether S_STD scales the transfer function for transitory signals without altering it. To evaluate the input/output transfer function I used 500 milliseconds test-trains of stimuli delivered at frequencies between 10 and 300Hz to match usual PC signals related to common movements, while the effect of S_STD was explored using protracted PC activation with different BFs prior the test-trains (Fig. 5A).

**Figure 5.**
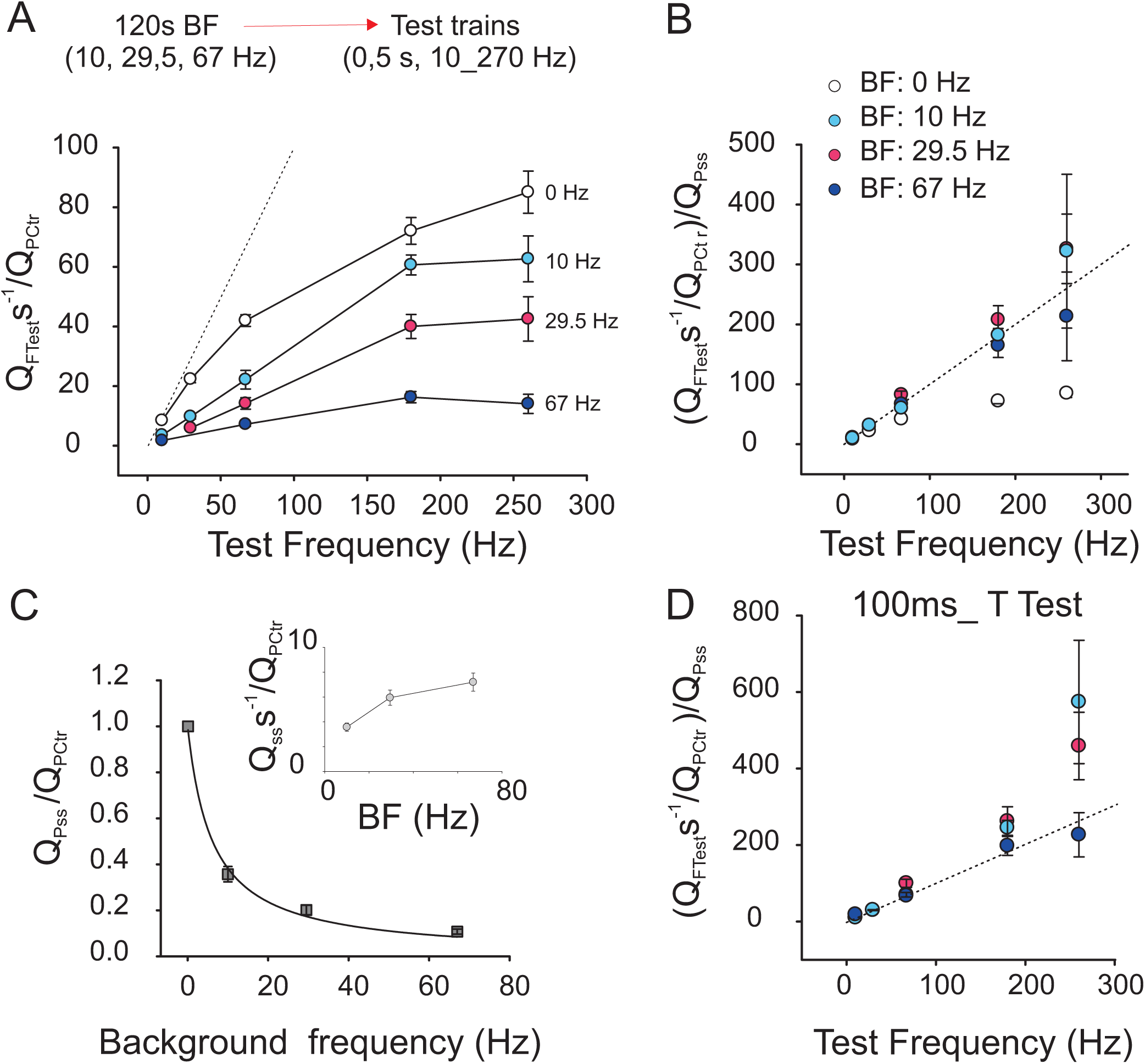
S_STD scales the PC_DCN transfer function of changes in PC rate. (A) Top: Stimulation paradigm. Bottom: plot of total charge transfer amplitude as a function of the test frequency for the 500 ms test-trains of stimuli applied from rest (0Hz), or after 2minutes of sustained stimulation at 10, 29.5 or 67 Hz (background frequency, BF). The responses were normalized to control IPSCs (dashed line: unity line). (B) Same as (A) but normalized to the charge transferred by IPSCs at steady state (IPSC_S_STDss_). The responses to test stimuli applied from rest, 0 Hz, in empty black. Note the plots overlap close to the unity line. Linear fits for the test frequency range 10-200 Hz (for clarity not shown): For BF 10 Hz: slope 1.0 ± 0.04, R^2^: 0.99; BF 29.5 Hz: slope 1.1 ± 0.06, R^2^: 0.99; BF 67 Hz: slope 0.9 ± 0.03, R^2^: 0.99. Dashed line=unity line. (C). Average normalized charge transfer of IPSC_S_STDss_ as a function of BF (n=25). The curve depicts the fit with a rational function (n=25, R^2^=0.999). Inset, depicts the relationship between the measured charge transfer at steady state per time unit IPSCs and the BF, normalized to control (Note different scale). (D) Same as (B) but for the first 100 ms of the responses to 500ms test trains.

I found the relationship between the average test-charge transfer amplitude and the test-frequency obtained from these experiments was approximately linear over the physiological range of PC rates (10 to 200 Hz) with slopes or gains variable and depending on the preceding BF. This is in agreement with a scaling effect and suggests the S_STD mechanism is independent of the one supporting frequency invariant transmission. In contrast, the transfer function curve obtained from rest was a saturating type of curve consistent with the predominantly fast depression of early events, as will be further discussed (0 Hz, black empty circles). Moreover, I explored whether the gains of the transfer functions for the test trains are only dependent on the preceding synaptic strength by renormalizing the data by the charge transferred by the corresponding IPSCs at steady state (IPSCs_S_STDSS_). The renormalization, in addition, should deduct potential biases induced by different levels of depression exhibited by different neurons (e.g. Figure 1D). Indeed, the results of this transformation (Figure 5B) show that, over the range 10 to 200 Hz, the slopes corresponding to different BFs coincide with the unity line, consistent with the notion that S_STD controls the gain according to the preceding level of depression without altering the linear transfer function (Figure 5B). This, together with recent reports of linear transfer function of PC_DCN synapses explored with short trains, when S_STD is not yet expressed ^10,11^, suggests that S_STD scales by a constant factor the transient responses without interfering with the mechanisms underlying linear encoding.

Next, I investigated the frequency dependence of IPSCs_STDSS_ to get an insight on the frequency dependence of S_STD itself. The average charge transferred by single IPSCs_STDSS_ as a function of preceding BF, normalized to control IPSCs, displayed a non-linear decrease with increasing BFs (Figure 4C, black line: fit using a rational function). This feature has been indicated as important for the re-scaling of responses to changes in rate (Abbott, 1997), as discussed later. Moreover and congruently, the measured total charge transferred at steady state, expressed per time unit and normalized to control IPSCs, increased non-linearly with BF displaying a shallow curve (Figure 5C inset, compare with Figure 5A), indicative of low sensitivity to BFs at steady state.

The switches to higher stimulation frequencies from steady state resulted in relative transient facilitation of PC_DCN_IPSCs amplitude (e.g. Figure 3B), in agreement with previous evidence of facilitation at these synapses from depressed states ^10–12^. How is the transfer of signals influenced by the presence facilitation? The analysis of the charge transferred during the first 100 ms of the test trains (Fig. 5D), when relative facilitation predominates, normalized to IPSCs_STDSS_ as a function of BF showed again an approximately linear function within functional range but with slopes slightly higher than one. This property could be useful for preferential transfer of brief transient signals, or the onset of longer ones. In addition, for the higher test-frequencies (260 Hz), the plots tended to deviated from linearity with diverging directions depending on the preceding BF: after 10 or 29.5 Hz, the responses were often supra-linear, although variable (note the high SEM), in line with the idea that similar type of synapses differ mainly by the degree of facilitation present ^38^. In brief, facilitation at PC_DCN synapses may not only support linear encoding by counteracting frequency dependent fast depression ^10,11^, but have a role in differentiating the length of signals, boosting the transfer of brief or the onset PC rate change signals. In addition, the data suggested that the ability to transfer brief and high frequency signals, e.g. complex spikes, might depend on the preceding BF.

Overall, the results suggest that S_STD de-emphasizes differences in steady-state PC rates, while enabling for a time window fitting most common motor behavior, a scaled linear relationship between the behavior-driven PC rates and the PC_DCN synaptic output. This opens the question of how S_STD, co-expressed with other forms of short-term plasticity, operates and interacts to determine this result. To answer this question, first, I explored the basis of S_STD and next, I used this information to implement a computational model featuring a slow depression component.

### Input independence and pre-synaptic location of S_STD mechanism

To investigate S_STD input-independence and because activation of two independent sets of PC axons could not be achieved under the used recording conditions, I combined opto-genetic (OG) and electrical stimulation (ES) in slices from mice expressing channelrhodopsin in PCs under the L7 promoter (L7-ChR2-eYFP ^39^) (Fig. 6A-B). The rational followed was, if depression was input-independent, tonic electrical stimulation should depress synaptic transmission at electrically stimulated terminals and spare those activated only by light. Indeed, the results illustrated in Fig. 6A-B show that after 2 minutes of repetitive electrical stimulation at 67 Hz, the depression of a larger OG-IPSCs (larger number of terminals involved) than a ES-IPSCs was significantly less, supporting the idea that depression is input specific.

**Figure 6.**
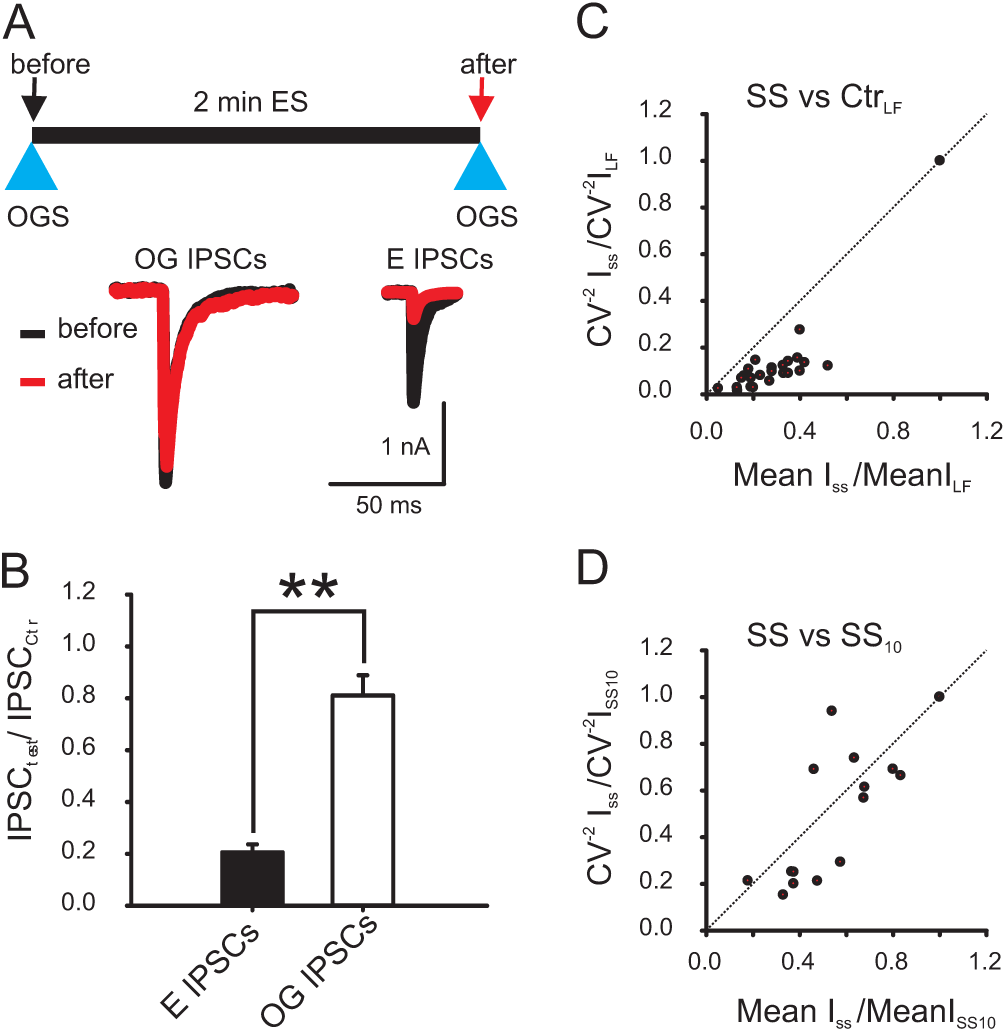
Input specificity and presynaptic origin of S_STD mechanism. (A) Top: depicts the experimental paradigm. Bottom: depicts representative examples of OG-IPSCs and ES-IPSCs before (control, black) and after (test, red) 2 minutes of PC electrical stimulation at 67 Hz.. (B) Plot of average OG- and ES-IPSCs amplitude after 2 minutes PC-ES at 67 Hz, normalized to their corresponding controls (IPSCs induced after 2 minutes without ES), (0.81±0.078 vs. 0.21±0.03; significance assessed with t-test, P<0.001, n=8). (C) Plot depicting changes in IPSCs CV^-2^ and their corresponding mean amplitudes at steady state after 2 min. stimulation with 10, 29.5 or 67 Hz normalized to control IPSCs evoked at low frequency (0.1-0.05 Hz) (n=8). (D), idem but for IPSCs depressed using BF 29.5 or 67 Hz normalized to values using BF 10 Hz (n= 7). Dashed line is unity line. Fit of data using a linear function, R^2^:0.78, slope 1.07±0.13, n=7.

To gain further insight into the basis of the S_STD, the locus of synaptic depression was explored using the inverse squared coefficient of variation (CV^-2^) compared to the changes in respective means. This method differentiate pre-from post-synaptic mediated changes in synaptic transmission, due to insensitiveness to changes caused mainly by post-synaptic mechanisms ^40,41^, as confirmed earlier at PC_DCN synapses ^12^. The CV^-2^ of IPSCs depressed to steady-state, normalized to control IPSCs evoked using low frequency (0.1-0.05Hz), decreased more than the normalized mean, a result compatible with a pre-synaptic locus of plasticity and moreover, assuming a binomial model of release ^41^, with a reduction in the probability of neurotransmitter release (Figure 6C). Next, to explore the basis of S_STD we compared IPSCs depressed to steady state using different BFs, normalized to the corresponding 10 Hz values (Figure 6D). The decrease in normalized CV^-2^ indicated a pre-synaptic mechanism. In this case, the CV^-2^ decreased in a similar proportion to the means, in agreement with a change in binomial “N”^41^, as expected e.g. from a depletion mechanism. These results suggest involvement of different presynaptic mechanisms: an early one predominantly dependent on a change in release probability (for synapses challenged from rest or low activation rate) and a different one involving a change in N (when comparing synapses depressed to stead state with different BFs). The change in release probability is in agreement with idea that at rest vesicles or sites show heterogeneous release properties ^42,43^ leading to early fast depletion of vesicles/sites displaying high release probability and slow rate of recovery. The use of this pool of vesicles could be at the basis for the change from a saturating to a linear transfer function (Fig. 5A, B).

Beyond these considerations, these results taken together are in agreement with the idea that S_STD is pre-synaptic and input specific.

### A model with a slow depression component explains well the experimental results

To gain insight onto how the interaction of different synaptic components explains the present results I used computer simulations. First, I tested a version of previously published models which features two different pools of ready to be released vesicles (RRPA and RRPB) ^43^. A version of this model including facilitation of release has been shown to explain well PC_DCN responses to hundreds of stimuli ^10,11^, and further agrees with the present CV^2^ results (See Methods for details). Briefly, the RRPs of this model (A and B) differ in release probability (Pr), presence of facilitation (F) and rate of recovery after release (Tau_r_). In this version, facilitation results from increased release probability and fast depression from depletion of RRPs (hereafter D+F model, Fig. 7 scheme; see Methods for details). The D+F model explained well the present PC_DCN responses to the first one hundred stimuli of trains with frequencies 10, 29.5 and 67 Hz (same data as in Fig. 1), showing frequency invariance and approximately linear encoding explained by the counteracting facilitation and depression, confirming previous results ^10^.

**Figure 7.**
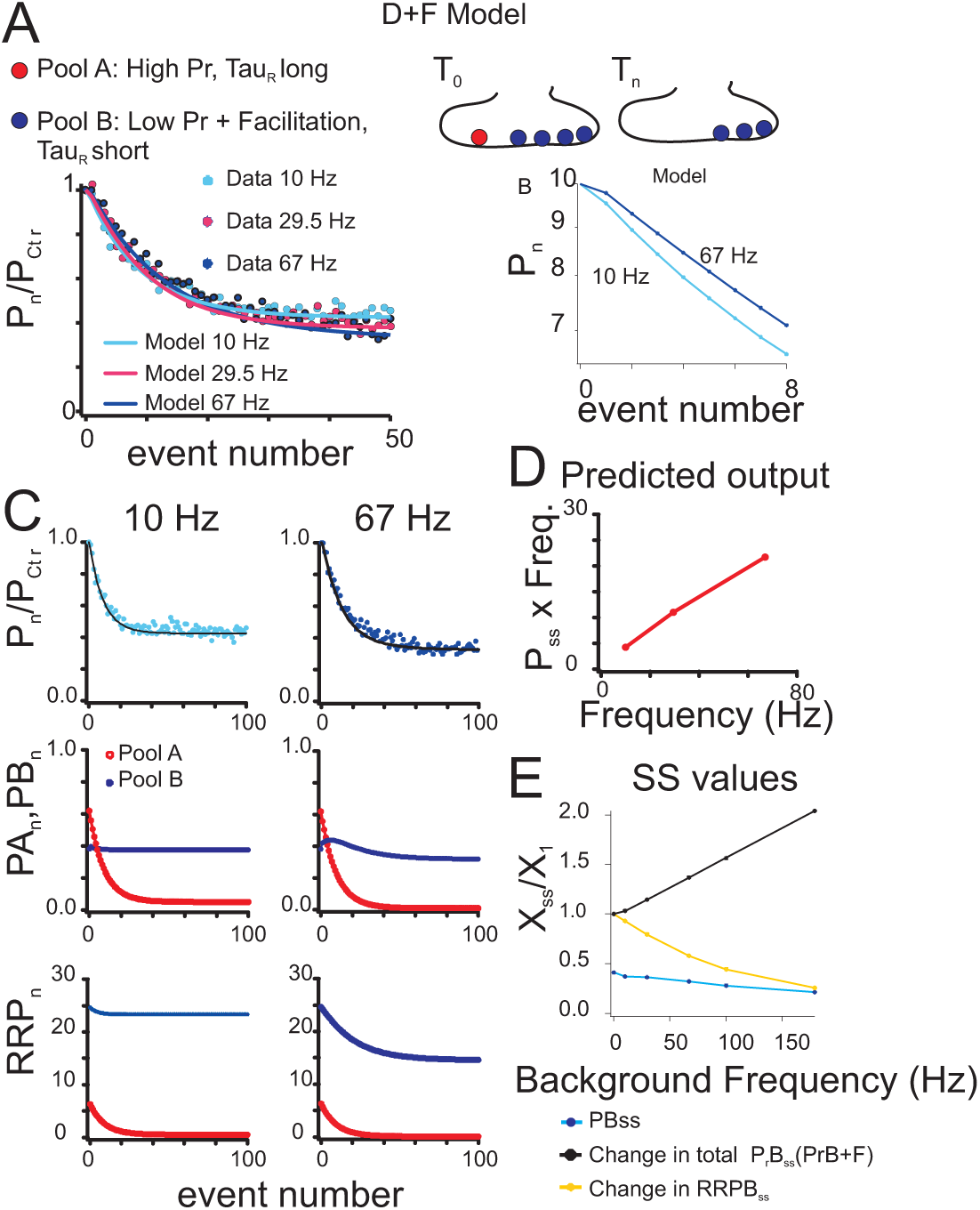
Experimental and simulated responses of a two-pool and facilitation model of PC_DCN synapses to different stimulation frequencies. (A) Top, D+F model scheme. Bottom, the predicted output of a two-pools and facilitation model (D+F model) explains the experimental responses to the first 100 events of stimulation trains at 10, 29.5 and 67 Hz (same data as in Fig. 1). (B) Detail of the model responses to the first 8 events of the trains at 10 and 67 Hz using a semi-logarithmic plot to illustrate slower decaying rates with higher stimulating frequencies. (C) Left and right plots (for 10 Hz and 67 Hz stimulation rates, respectively) illustrate from top to bottom, the data and model output, the predicted IPSC amplitudes for Pools A and B (PA, PB, red and blue) and the size of the ready to be released A and B pools (RRPA, RRPB, red and blue respectively). (D The relationship of the predicted synaptic output at steady state (estimated by the product of the predicted IPSC amplitude and the stimulation frequency) against the stimulation frequency is almost linear. (E) Summary of results predicted by the D+F model at steady state as a function of the stimulation frequency: the predicted output of the model (PB, blue circles), the total probability of release of the Pool B normalized to event 1 (P_r_B+facilitation, black circles), and the change in RRPB (yellow circles), normalized to event 1. (RRPA_0_=7, RRPB_0_=25, PrA=0.098, PrB=0.017, TaurA=12,TaurB=0.5, f1=0.0005, TauF1=0.007, f2=0.001, Tauf2=0.1).

However, this model could not explain the further decay in IPSC amplitude with protracted stimulation (Figure 8A), even when facilitation was not included. The mechanism of slow depression is debated, but it has been recently proposed that under intense or sustained synaptic activity the most sensitive bottleneck for the availability of vesicles for release is the number of release sites that after undergoing exocytosis are cleared and prepared to receive new vesicles ^44^. Considering (1), the latter idea (2), the frequency dependence of S_STD and (3), the S_STD dependence on the binomial value N suggested by present results (Fig. 6), I implemented a slow depression component by an activity-dependent decrease in active release sites of pool B (R_SitesB) recovering with fixed time constant (Fig. 8B and Methods). This component was used to extend the D+F model The extended model (hereafter, SD_RS model) was able to explain the changes in IPSC amplitude observed during 120s long stimulation trains (Fig. 8B).

**Figure 8.**
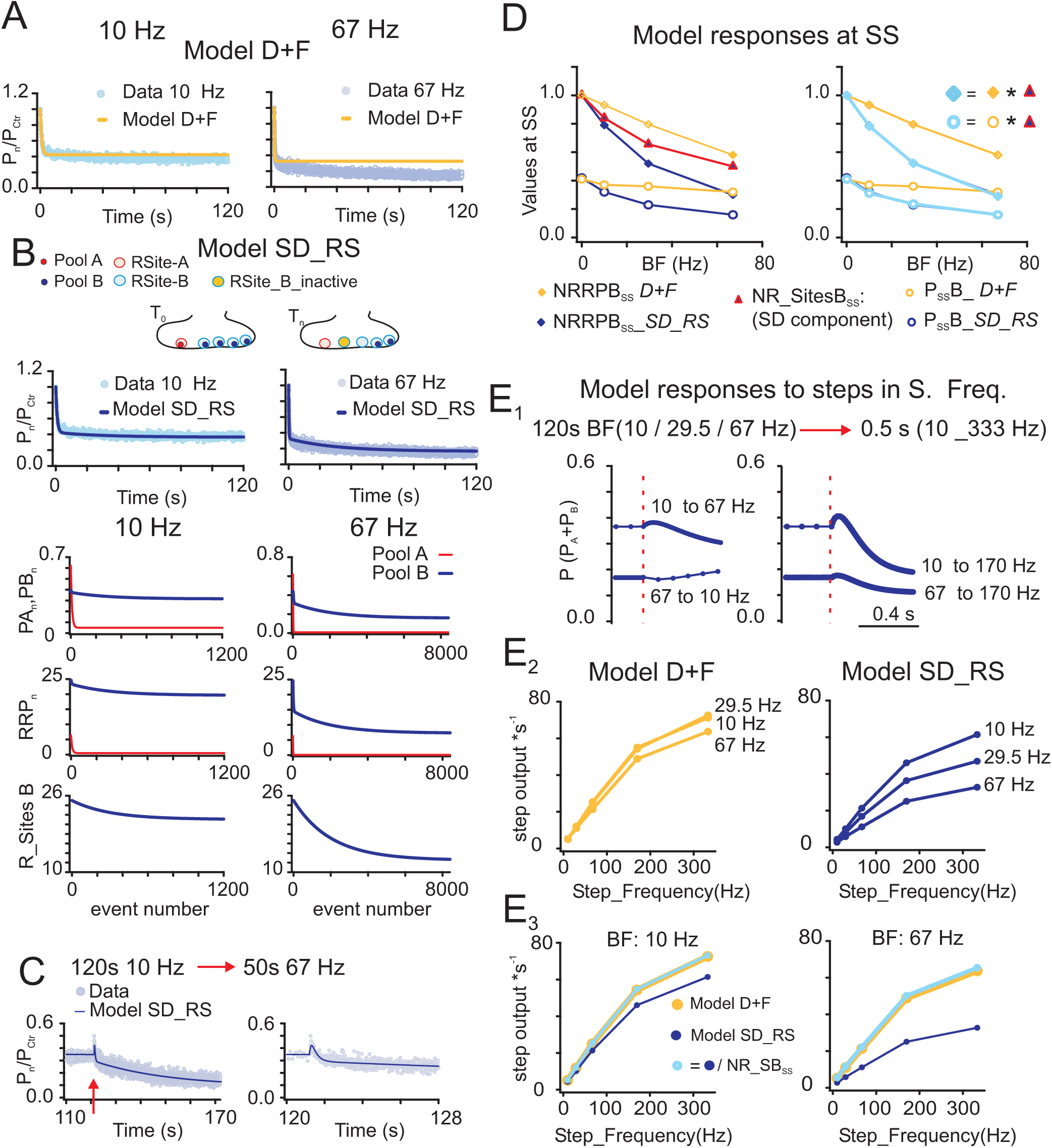
A two pool and facilitation model extended by a slow depression component (SD) explains experimental S_STD and resulting gain modulation of transient responses at PC_DCN synapses. (A) Predictions from a two-pool-and-facilitation model (D+F model) (yellow line, same fitting parameters as in Fig. 7) fail to explain S_STD (experimental data: blue and violet circles for 10 and 67 Hz, respectively, same data as in Fig. 1). (B) Top, instead, both, early and late experimental responses were explained by a model featuring a slow depression (SD) component schematized on top (SD-RS model: dark blue lines, see main text for details, data as in A). In lower rows from top to bottom Pool A (red) and B (blue): the predicted IPSC amplitudes of pools A and B (PA, PB), size of (filled) ready-to-be-released A and B pools (RRPA, RRPB), and the number of active release sites of pool B (R_SitesB). (C) Experimental (light blue) and SD-RS model predictions (dark blue line) to a sustained change in stimulation frequency (120s at 10 Hz followed by 50s at 67 Hz). Right plot, detail in expanded time scale, around the switch time. Data from a different set of neurons than A and B, n=6, the estimated parameters are in the legend of Fig. 9. (D) Left panel: predicted normalized RRPB (NRRPB, filled squares) and PB (open circles) values by the D+F (yellow) and SD_RS (dark blue) models and the slow depression component (red triangles, fractional change in R_SitesB of the RS-SD model) at steady state as a function of the background frequency (BF) (from the experiments illustrated in A and B). Right, a multiplicative effect of the SD component explains the differences between the models: the light blue curves are the product of the D+F curves (yellow) times the corresponding fractional R_SitesB (shown on the left plot in red); the dark blue traces (SD_RS model predictions) are occluded by the light blue curves. (E_1_) Model predictions to steps of change in stimulating frequency: Top: stimulation paradigm. Bottom: examples of SD_RS model responses (red dashed line indicates beginning of the step). (E_2_) Integral of predicted IPSCs during the 0.5 s steps (per sec), as a function of the step frequency, for different BFs (labeled on the right of each trace) and for the D+F and SD_RS models (left and right plots respectively). (E_3_) Multiplicative effect of SD: predicted input/output functions by the D+F (yellow) and SD-RS (dark blue) models as in E_2_, for 10 and 67 Hz BF (left and right plots respectively); the light blue traces were obtained by dividing the output of the SD-RS model (dark blue) by the corresponding fraction of R_SitesB available *before* the frequency step. (SD-RS: RRPA_0_=7, RRPB_0_= 25, PrA=0.098, PrB=0.01, TauRA=12, TauRB=1.55, f1=0.0005, TauF1=0.007, f2=0.0024, Ttauf2=0.1, ARSB=0.74, F_RS= 29, TauRSr= 30, apply to all panels except Fig. 6C, see details in main text).

Importantly, this model explained as well the results from a different set of experiments using a switch in the protracted stimulation frequency within the same trial, i.e., from 10 to 67 Hz (Fig. 8C, Fig. 9). These experimental results confirmed those obtained using cell-attached recordings (Figs. 4C-D): after a switch in BF it took several tens of seconds to reach a new steady state. In the model correlated with the slow change in “active release sites” after the switch (RSites B, Fig. 9D inset). Note the similarity in the time course of the amplitudes of the experimental and predicted responses: first a relative facilitation of IPSCs amplitude after the switch, which recovers and afterwards slowly re-adapt up to a new steady state level.

**Figure 9.**
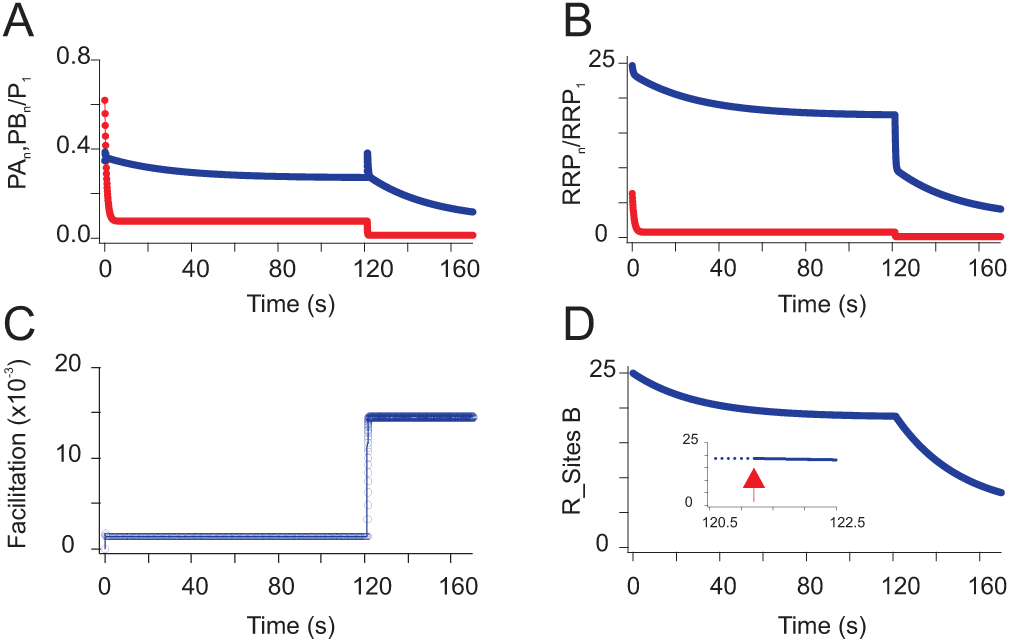
Predicted responses of the SD-RS model to a change in sustained stimulation frequency from 10 Hz (120s) to 67 Hz (50s). (A) PA and PB (red and blue, respectively, as in Fig.S2). (B) RRPA and RRPB (red and blue, respectively, as in Fig.S2). (C) Total facilitation for pool B. (D) Number of release sites active (receptive and filled, R_SitesB). Inset, depicts in expanded scale around the switch time (red arrow) the very low rate of change in R_SitesB. (RRPA_0_=7, RRPB_0_=25, PrA=0.098, PrB=0.01, TauRA=12, TauRB=1.55, f1=0.0005, TauF1=0.007, f2=0.0024, Tauf2=0.1, ARSB=0.74, F_RS= 29, TauRSr= 30).

The comparison of steady state values of RRPB size (diamonds) and the output of pool B (PB, circles) predicted by the SD_RS (blue) and the D+F (yellow) models—which differ only by the presence or absence of the slow-depression component—highlights the effect of the latter (Figure 8D, left panel). The plot illustrates too, the frequency dependent change in the slow-depression component represented by the fractional size of active release sites of pool B (R_SitesB, red triangles).

The scaling effect of S_STD (Figure 5B), suggests a multiplicative relationship of the slow depression component. The here implemented inactivation of sites does not interfere with the properties of the remaining active ones, predicting a scaling effect. Indeed, the product of the RRPB or the PB values of the D+F model times the slow-depression component return the RRPB and PB values of the SD_RS model respectively (Fig. 8D right, the respective products are shown in light blue), graphically demonstrating a multiplicative operation of the slow depression component at steady state, which was able to explain the effects of sustained PC activation. Noteworthy, similar results emerged for analysis of the total output (pool-A and -B) caused by minimal contribution of Pool-A at steady state (Fig. 10). Therefore, the extended model explains both the changes in amplitude and the time course of these changes during protracted stimulation and after a switch in BF.

**Figure 10.**
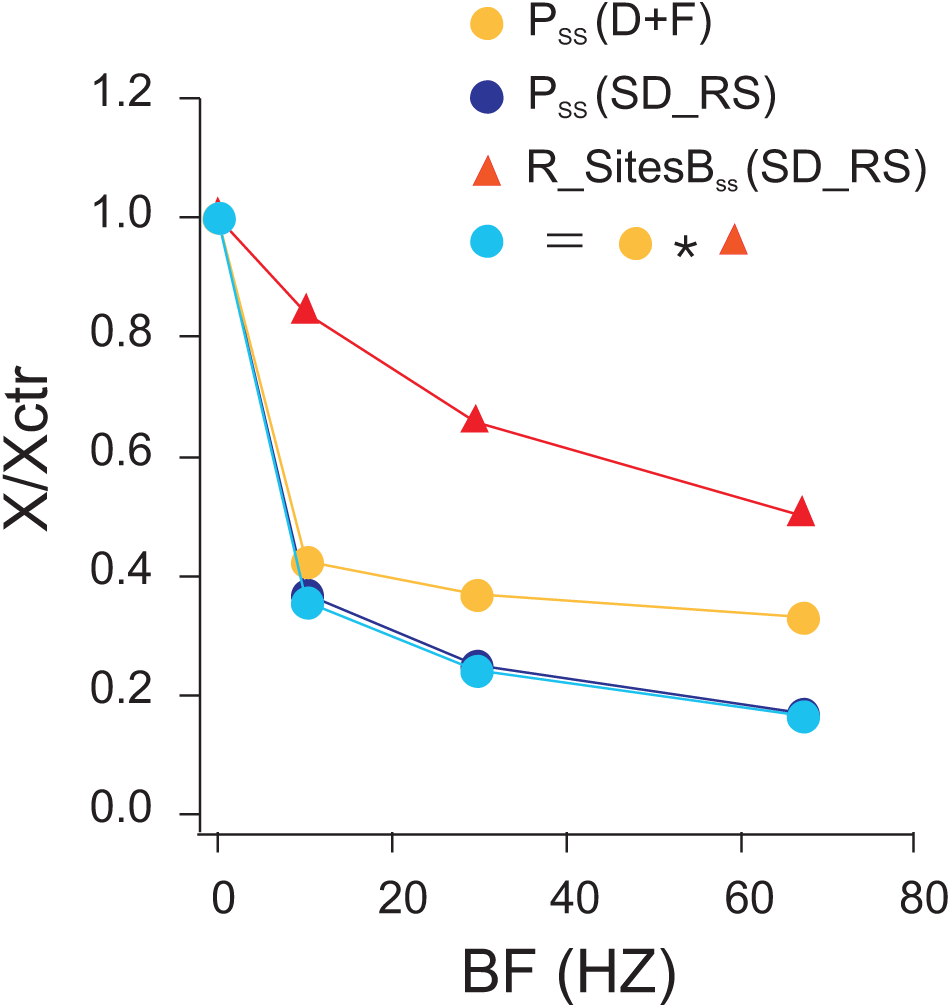
A multiplicative operation of the SD component (R_SitesB) explain he differences in the predicted steady state output of the D+F and SD-RS models as a function of BFs. The plot illustrates the IPSC amplitudes (PoolA+PoolB) at steady state predicted by the D+F (P_ss_, yellow) and SD_RS (P_ss_, dark blue) models, and the normalized values of the available release sites of the RS_SD model (R_SitesB, red). The result of multiplying the P_ss_ of the D+F model times the corresponding R_SitesB of the SD_RS model is shown as a function of the background stimulating frequency (light blue).

Therefore, this model is likely to be able to account for the responses to changes in stimulating frequency. To test this possibility, I compared the prediction of both models to steps of changes in activation frequency (duration: 0.5s) induced after sustained activation, as applied experimentally to PC_DCN synapses (Figure 8E1). As expected, the integral of the predicted IPSCs evoked over the step duration (proxy for the charge transfer amplitude) by the D+F model increases almost linearly as a function of the step frequency up to 200 Hz but the slopes for different BFs are almost coincident in contrast to the experimental results (Figure 8E2-left panels vs. Fig. 5A). Instead, the predictions by the RS-SD model (Figure 8E2-right) show similar linear increases in output but clearly distinct slopes for different BFs, indicating that the model captures the gain modulation for transitory changes in rate mediated by S_STD. Moreover, dividing the values obtained with the SD_RS model by the number of release sites present at steady state before the step returned the values of the D+F model (Figure 8E3), indicating that the multiplicative effect of this slow depression component of Pool B was sufficient to explains the effect. Importantly, the calculation used the number of release sites available just before the step, indicating that given the slow kinetics of this component (Fig. 9D), its state determines the gain for post-ceding sub-second-to-few seconds periods. The fast STP components (facilitation and depletion) led to opposing effects during the step, (as exemplified in Fig. 9, Turecek, 2016). Therefore, together the slow dynamics of the slow depression component and balanced fast facilitation and depression, explain the preserved gain over the first second after the change in PC rate.

Summarizing, a model of synaptic transmission including a slow depression factor explains well the results to PC_DCN protracted stimulation as well as the responses to changes in PC activity. Moreover, these results demonstrate that an activity dependent change in release sites is a plausible molecular mechanism for slow depression at PC_DCN synapses and consequent gain control.

## DISCUSSION

This study unveils a novel slow-gain-control mechanism at PC_DCNs, which reconciles linear encoding with adaptation to background-activity. The leading finding was the identification of a previously unrecognized PC-DCNs frequency-dependent slow short-term depression (S_STD), evolving over tens of seconds, i.e., orders of magnitude slower than fast STD forms, allowing PC_DCN adaptation to background activity. I show that after changes in PC_DCNs activation-rate (used to mimic the behavior-related PC signals embedded in the sustained activity) the synaptic gain remained stable over second-long time windows, supporting scaled linear encoding of PC rates in the timescale of common movements (reaches, steps, etc.). Thus, S_STD enables the assignment of different functions to different timescales. The results from experimental and simulation analyses suggested that the independent and multiplicative relationship of two synaptic mechanisms, one explaining slow depression and the other frequency-invariant transmission, could account for S_STD-gain-control. Furthermore, evidence from opto-genetic experiments, statistical analysis and computer simulation pointed to a presynaptic, input specific and possibly activity-dependent decrease in release-sites as the ground for S_STD and derived computations. I argue this novel slow-gain-control mechanism may first, explain how PCs can effectively transfer linearly encoded behavior-related signals despite disparate background-rates, and second, provide an additional channel for fine-tuning each PC output at its last stage. Furthermore, because slow movements apparently result from the integration of shorter sub-movements ^45^, S_STD-gain-control might serve to linearly transfer the corresponding chunks of PC information. Finally, because sustained neuronal activity is inherent to numerous neural processes, S_STD-gain-control might be fundamental to other functions in different brain areas.

The use of protracted PC_DCN activation has been key in the identification of S_STD, which was a robust finding, revealed using regular and irregular activation patterns in preparations from young or adult animals and under conditions alike those found *in situ* (i.e., temperature, calcium concentration). Furthermore, the effect of S_STD modulation on DCN responses was readily assessable using whole-cell patch (charge transfer) and cell-attached DCN recordings (changes in firing rate), confirming its functional significance. As the use of protracted synaptic activation tests is rare, the prevalence of slow depression at other synapses is unclear. However, evidence for S_STD was first recognized at the neuromuscular junction ^46,47^ and subsequently in hippocampal cell cultures, hippocampal slices and the auditory pathway ^48–52^. However, slow STDs probably only arise under sustained activation, a condition clearly met at PC_DCNs ^16,53–56^, but apparently not at hippocampal synapses ^57^.

The mechanism/s of S_STD is debated, with proposed mechanisms including depletion of a reserve-pool ^47,58^, “fatigue” in the replenishment of the readily releasable pool of vesicles (RRP) ^59^ and decreased release probability ^51^. The latter mechanism seems unlikely for PC_DCNs given the CV^-2^ analysis suggesting decreased “N”, but see ^60^. The same results speak against a decreased quantal size, e.g. by incomplete vesicle refilling ^61^. Furthermore, GABA_B_-dependent slow gain-control ^62^ is unlikely the cause for PC_DCN-S_STD as the time course differs, and PC_DCN-S_STD modulation persists under GABA_B_ antagonists, in line with previous results ^12^. Recently the availability of “receptive” release-sites has been proposed to be the principal bottleneck for the maintenance of intense sustained synaptic transmission ^44^, suggested by observations of rapid reduced synaptic release under unbalanced exocytosis and endocytosis, faster than expected from a failure in vesicle recycling ^63,64^. The exact mechanism limiting release-sites availability is not clear (Rev. in ^65^), but could explain S_STD. In fact, changes in release sites was proposed earlier to explain a slow phase of recovery from intense activity ^49^. However, even if this mechanism results from a synaptic limitation, it must not be universally obligatory ^66^, as slow forms of synaptic enhancement ^67–69^ and/or different structural synaptic properties could mitigate this limitation ^65^.

In the present study, I implemented “release-site availability” as a component of a previously published synaptic transmission model explaining frequency-invariance on shorter time scales by balanced fast-short-term plasticity(f-STP) ^10^. The success of this extended model in predicting the experimental S_STD of PC-DCN IPSCs strongly favored “release-site availability” as PC_DCNs S_STD mechanism.

Furthermore, the extended model accounted for the responses to changes in PC_DCNs activation rate. Indeed, the experimental results showed that the synaptic gain remained stable within the first hundreds of milliseconds to seconds after a change in rate, enabling for each gain-level over this time window scaled linear encoding of the new PC rates. This observation suggested that the process underlying S_STD is different and independent from the one supporting frequency-invariance; otherwise, induction of S_STD could have altered the linear encoding, e.g. by unbalanced fast-facilitation and depression. An activity-dependent decrease in active-release-sites does not (necessarily) influence the properties of the remaining sites and thus is compatible with an independent mechanism further arguing for this as a mechanism of S-STD. The model accounted not only for the values at steady state but also for the time course of changes in IPSCs amplitudes evoked by different protocols (Figs. 8C and 9). Although the rate at which single vesicle-depleted release-sites recover might be activity-dependent ^70^, in the present model this rate was kept constant although the total vesicle-recover rate may vary due to activity-dependent changes in the total number of “accepting release-sites”. Nevertheless, enhanced vesicle recruitment could be one possible explanation for theobservation of supralinear responses using very high and brief changes in rate (Fig. 5D). Overall, this study supports a decrease in “accepting release-sites” as a plausible mechanism for S_STD and (together with fast frequency-invariant synaptic transmission) the derived slow-gain-control mechanism described here. Nonetheless, the molecular identity behind the S_STD phenomenon could correspond to an equivalent process, warranting further studies.

What functional consequences arise from PC_DCN-S_STD?

First, the mere presence of S_STD and the resulting shallow relationship between the background-stimulation frequency (BF) and the charge transfer amplitude at steady state (Fig. 6C, inset) suggests S_STD de-emphasizes the effect of PC background-activity in behaving animals. Thus, S_STD could partly explain previous findings of little sensitivity of DCN firing responses to sustained changes PC_BF^29^. The decreased sensitivity to steady PC rates could be useful for maintaining the PC output, which is inhibitory, within the working firing range of DCNs avoiding transmission failure ^71^.

Second, S-STD gain-control could explain how PC_DCNs could effectively convey linearly encoded behavior-driven PC signals despite PCs diverse and variable background-activity rates. Fast-STD dynamic gain-control mechanism was proposed to enhance the sensitivity of postsynaptic neurons to changes in their input rates, partly because at steady state the synaptic strength is approximately inversely proportional to the activation rate determining that equal fractional changes in rate produce equal changes in output, independently of absolute rates, a “Weber-Fechner law” effect, ^3^. The frequency-dependence of PC_DCN-IPSCs at steady state follows approximately a rational function (Fig. 5C, i.e., ca. inversely proportional to BF), suggesting similar property for S_STD, but effective over longer time windows. Indeed, similar fractional increases in PC rate resulted in similar outputs for different BFs (Fig. 5A, 10 to 30 Hz and 67 to 180 Hz).

Notably, this property could be useful for rate coding, but also for transferring the timing of synchronized (complex or simple) spikes, or firing pauses ^37,72–75^, or for graded signal inversion ^71,76^. Interestingly, facilitation supported higher sensitivity to brief changes in rate (or the onset of longer ones, Fig. 5D) suggesting the code might be time-dependent.

Finally, intriguingly, the changes in gain without changes in the transfer function of mimicked behavior-related PC signals induced by background-activity through S_STD is reminiscent, at synaptic level, of the gain-modulation of neuronal population responses ^77^, a computation proposed to account for non-linear combination of signals ^78^. PCs sustained firing and PC behavior-related signals converge at single synapses but each may represent different information and consequently, change independently. Thus, modulation of PC background-activity (via cerebellar long-term plasticity or context-dependent changes in afferent activity) without corresponding scaling of behavior-driven PC signals could modify PC output gain, in a similar way as shown here (e.g., Fig. 5A, changes from 10 vs. 67 Hz to 260 Hz). Therefore, these results suggest S-STD gain control could provide an alternative channel for flexibly tuning each PCs output at the last cerebellar stage ^79^, and point to a novel role for PCs background-activity control in cerebellar function.

## REFERENCES

1. Silver, R.A. Neuronal arithmetic. Nat Rev Neurosci 11, 474–489 (2010).

2. Whitmire, C.J. & Stanley, G.B. Rapid Sensory Adaptation Redux: A Circuit Perspective. Neuron 92, 298–315 (2016).

3. Abbott, L.F., Varela, J.A., Sen, K. & Nelson, S.B. Synaptic depression and cortical gain control. Science 275, 220–224 (1997).

4. Tsodyks, M.V. & Markram, H. The neural code between neocortical pyramidal neurons depends on neurotransmitter release probability. Proc. Natl. Acad. Sci. U. S. A 94, 719–723 (1997).

5. Zucker, R.S. & Regehr, W.G. Short-term synaptic plasticity. Annu. Rev. Physiol 64, 355–405 (2002).

6. Herzfeld, D.J., Kojima, Y., Soetedjo, R. & Shadmehr, R. Encoding of action by the Purkinje cells of the cerebellum. Nature 526, 439–442 (2015).

7. Ten Brinke, M.M., et al. Dynamic modulation of activity in cerebellar nuclei neurons during pavlovian eyeblink conditioning in mice. Elife 6 (2017).

8. Romano, V., et al. Potentiation of cerebellar Purkinje cells facilitates whisker reflex adaptation through increased simple spike activity. Elife 7 (2018).

9. Catz, N., Dicke, P.W. & Thier, P. Cerebellar-dependent motor learning is based on pruning a Purkinje cell population response. Proc Natl Acad Sci U S A 105, 7309–7314 (2008).

10. Turecek, J., Jackman, S.L. & Regehr, W.G. Synaptic Specializations Support Frequency-Independent Purkinje Cell Output from the Cerebellar Cortex. Cell Rep 17, 3256–3268 (2016).

11. Turecek, J., Jackman, S.L. & Regehr, W.G. Synaptotagmin 7 confers frequency invariance onto specialized depressing synapses. Nature (2017).

12. Pedroarena, C.M. & Schwarz, C. Efficacy and short-term plasticity at GABAergic synapses between Purkinje and cerebellar nuclei neurons. J Neurophysiol 89, 704–715 (2003).

13. Arenz, A., Silver, R.A., Schaefer, A.T. & Margrie, T.W. The Contribution of Single Synapses to Sensory Representation in Vivo. Science 321, 977 (2008).

14. Jelitai, M., Puggioni, P., Ishikawa, T., Rinaldi, A. & Duguid, I. Dendritic excitation–inhibition balance shapes cerebellar output during motor behaviour. Nature Communications 7, 13722 (2016).

15. Chen, S., Augustine, G.J. & Chadderton, P. The cerebellum linearly encodes whisker position during voluntary movement. eLife 5, e10509 (2016).

16. Hong, S., et al. Multiplexed coding by cerebellar Purkinje neurons. eLife 5 (2016).

17. Dugué, G.P., Tihy, M., Gourévitch, B. & Léna, C. Cerebellar re-encoding of self-generated head movements. eLife 6, e26179 (2017).

18. Halverson, H.E., Khilkevich, A. & Mauk, M.D. Relating cerebellar purkinje cell activity to the timing and amplitude of conditioned eyelid responses. J Neurosci 35, 7813–7832 (2015).

19. Thach, W.T. Discharge of Purkinje and cerebellar nuclear neurons during rapidly alternating arm movements in the monkey. J Neurophysiol 31, 785–797 (1968).

20. Hobson, J.A. & McCarley, R.W. Spontaneous discharge rates of cat cerebellar Purkinje cells in sleep and waking. Electroencephalogr. Clin. Neurophysiol 33, 457–469 (1972).

21. De Zeeuw, C.I., Wylie, D.R., Stahl, J.S. & Simpson, J.I. Phase relations of Purkinje cells in the rabbit flocculus during compensatory eye movements. J Neurophysiol 74, 2051–2064 (1995).

22. Bryant, J.L., Roy, S. & Heck, D.H. A technique for stereotaxic recordings of neuronal activity in awake, head-restrained mice. J Neurosci Methods 178, 75–79 (2009).

23. Zhou, H., et al. Cerebellar modules operate at different frequencies. eLife 3, e02536 (2014).

24. Laurens, J. & Angelaki, D.E. A unified internal model theory to resolve the paradox of active versus passive self-motion sensation. Elife 6 (2017).

25. Xiao, J., et al. Systematic Regional Variations in Purkinje Cell Spiking Patterns. PLOS ONE 9, e105633 (2014).

26. Rinaldo, L. & Hansel, C. Ataxias and cerebellar dysfunction: involvement of synaptic plasticity deficits? Funct Neurol 25, 135–139 (2010).

27. Tsai, P.T., et al. Autistic-like behaviour and cerebellar dysfunction in Purkinje cell Tsc1 mutant mice. Nature 488, 647–651 (2012).

28. Ozden, I., Dombeck, D.A., Hoogland, T.M., Tank, D.W. & Wang, S.S. Widespread state-dependent shifts in cerebellar activity in locomoting mice. PLoS One 7, e42650 (2012).

29. Belmeguenai, A., et al. Intrinsic plasticity complements long-term potentiation in parallel fiber input gain control in cerebellar Purkinje cells. J Neurosci 30, 13630–13643 (2010).

30. Shim, H.G., et al. Long-Term Depression of Intrinsic Excitability Accompanied by Synaptic Depression in Cerebellar Purkinje Cells. J Neurosci 37, 5659–5669 (2017).

31. Gao, Z., van Beugen, B.J. & De Zeeuw, C.I. Distributed synergistic plasticity and cerebellar learning. Nat Rev Neurosci 13, 619–635 (2012).

32. Magleby, K.L. The effect of tetanic and post-tetanic potentiation on facilitation of transmitter release at the frog neuromuscular junction. J Physiol 234, 353–371 (1973).

33. MacLeod, K.M., Horiuchi, T.K. & Carr, C.E. A role for short-term synaptic facilitation and depression in the processing of intensity information in the auditory brain stem. J Neurophysiol 97, 2863–2874 (2007).

34. Doussau, F., et al. Frequency-dependent mobilization of heterogeneous pools of synaptic vesicles shapes presynaptic plasticity. Elife 6 (2017).

35. Uusisaari, M., Obata, K. & Knopfel, T. Morphological and Electrophysiological Properties of GABAergic and Non-GABAergic Cells in the Deep Cerebellar Nuclei. Journal of Neurophysiology 97, 901–911 (2007).

36. Telgkamp, P. & Raman, I.M. Depression of inhibitory synaptic transmission between Purkinje cells and neurons of the cerebellar nuclei. J Neurosci 22, 8447–8457 (2002).

37. Shin, S.L., et al. Regular patterns in cerebellar Purkinje cell simple spike trains. PLoS. One 2, e485 (2007).

38. Fekete, A., et al. Underpinning heterogeneity in synaptic transmission by presynaptic ensembles of distinct morphological modules. Nat Commun 10, 826 (2019).

39. Chaumont, J., et al. Clusters of cerebellar Purkinje cells control their afferent climbing fiber discharge. Proceedings of the National Academy of Sciences 110, 16223–16228 (2013).

40. Bekkers, J.M. & Stevens, C.F. Presynaptic mechanism for long-term potentation in the hippocampus. Nature 346, 724–729 (1990).

41. Malinow, R. & Tsien, R.W. Presynaptic enhancement shown by whole-cell recordings of long-term potentiation in hippocampal slices. Nature 346, 177–180 (1990).

42. Rosenmund, C., Clements, J.D. & Westbrook, G.L. Nonuniform probability of glutamate release at a hippocampal synapse. Science 262, 754–757 (1993).

43. Trommershauser, J., Schneggenburger, R., Zippelius, A. & Neher, E. Heterogeneous presynaptic release probabilities: functional relevance for short-term plasticity. Biophys. J 84, 1563–1579 (2003).

44. Neher, E. What is Rate-Limiting during Sustained Synaptic Activity: Vesicle Supply or the Availability of Release Sites. Front Synaptic Neurosci 2, 144 (2010).

45. Park, S.W., Marino, H., Charles, S.K., Sternad, D. & Hogan, N. Moving slowly is hard for humans: limitations of dynamic primitives. J Neurophysiol 118, 69–83 (2017).

46. Del Castillo, J. & Katz, B. Statistical factors involved in neuromuscular facilitation and depression. J Physiol 124, 574–585 (1954).

47. Elmqvist, D. & Quastel, D.M. A quantitative study of end-plate potentials in isolated human muscle. J. Physiol 178, 505–529 (1965).

48. Liu, G. & Tsien, R.W. Properties of synaptic transmission at single hippocampal synaptic boutons. Nature 375, 404–408 (1995).

49. Stevens, C.F. & Wesseling, J.F. Identification of a novel process limiting the rate of synaptic vesicle cycling at hippocampal synapses. Neuron 24, 1017–1028 (1999).

50. Garcia-Perez, E., Lo, D.C. & Wesseling, J.F. Kinetic isolation of a slowly recovering component of short-term depression during exhaustive use at excitatory hippocampal synapses. J Neurophysiol 100, 781–795 (2008).

51. Hennig, M.H., Postlethwaite, M., Forsythe, I.D. & Graham, B.P. Interactions between multiple sources of short-term plasticity during evoked and spontaneous activity at the rat calyx of Held. J. Physiol 586, 3129–3146 (2008).

52. Krächan, E.G., Fischer, A.U., Franke, J. & Friauf, E. Synaptic reliability and temporal precision are achieved via high quantal content and effective replenishment: auditory brainstem versus hippocampus. The Journal of Physiology 595, 839–864 (2017).

53. Schonewille, M., et al. Purkinje cells in awake behaving animals operate at the upstate membrane potential. Nat Neurosci 9, 459–461 (2006).

54. Norris, S.A., Greger, B., Hathaway, E.N. & Thach, W.T. Purkinje cell spike firing in the posterolateral cerebellum: correlation with visual stimulus, oculomotor response, and error feedback. J Neurophysiol 92, 1867–1879 (2004).

55. Sato, Y., Miura, A., Fushiki, H. & Kawasaki, T. Short-term modulation of cerebellar Purkinje cell activity after spontaneous climbing fiber input. Journal of Neurophysiology 68, 2051 (1992).

56. Cao, Y., Maran, S.K., Dhamala, M., Jaeger, D. & Heck, D.H. Behavior-related pauses in simple-spike activity of mouse Purkinje cells are linked to spike rate modulation. J Neurosci 32, 8678–8685 (2012).

57. Klyachko, V.A. & Stevens, C.F. Excitatory and feed-forward inhibitory hippocampal synapses work synergistically as an adaptive filter of natural spike trains. PLoS Biol 4, e207 (2006).

58. Richards, D.A., Guatimosim, C., Rizzoli, S.O. & Betz, W.J. Synaptic vesicle pools at the frog neuromuscular junction. Neuron 39, 529–541 (2003).

59. Gabriel, T., et al. A new kinetic framework for synaptic vesicle trafficking tested in synapsin knock-outs. J. Neurosci 31, 11563–11577 (2011).

60. Faber, D.S. & Korn, H. Applicability of the coefficient of variation method for analyzing synaptic plasticity. Biophys J 60, 1288–1294 (1991).

61. Bennett, M.V., Model, P.G. & Highstein, S.M. Stimulation-induced depletion of vesicles, fatigue of transmission and recovery processes at a vertebrate central synapse. Cold Spring Harb Symp Quant Biol 40, 25–35 (1976).

62. Magnusson, A.K., Park, T.J., Pecka, M., Grothe, B. & Koch, U. Retrograde GABA signaling adjusts sound localization by balancing excitation and inhibition in the brainstem. Neuron 59, 125–137 (2008).

63. Kawasaki, F., Hazen, M. & Ordway, R.W. Fast synaptic fatigue in shibire mutants reveals a rapid requirement for dynamin in synaptic vesicle membrane trafficking. Nat Neurosci 3, 859–860 (2000).

64. Hosoi, N., Holt, M. & Sakaba, T. Calcium dependence of exo- and endocytotic coupling at a glutamatergic synapse. Neuron 63, 216–229 (2009).

65. Byczkowicz, N., Ritzau-Jost, A., Delvendahl, I. & Hallermann, S. How to maintain active zone integrity during high-frequency transmission. Neurosci Res 127, 61–69 (2018).

66. Mayer, F., Albrecht, O., Dondzillo, A. & Klug, A. Glycinergic inhibition to the medial nucleus of the trapezoid body shows prominent facilitation and can sustain high levels of ongoing activity. J Neurophysiol 112, 2901–2915 (2014).

67. Xue, R., et al. Doc2-mediated superpriming supports synaptic augmentation. Proc Natl Acad Sci U S A 115, E5605–E5613 (2018).

68. Kandaswamy, U., Deng, P.Y., Stevens, C.F. & Klyachko, V.A. The role of presynaptic dynamics in processing of natural spike trains in hippocampal synapses. J Neurosci 30, 15904–15914 (2010).

69. Kalkstein, J.M. & Magleby, K.L. Augmentation increases vesicular release probability in the presence of masking depression at the frog neuromuscular junction. J Neurosci 24, 11391–11403 (2004).

70. Pan, B. & Zucker, R.S. A general model of synaptic transmission and short-term plasticity. Neuron 62, 539–554 (2009).

71. Pedroarena, C.M. Mechanisms supporting transfer of inhibitory signals into the spike output of spontaneously firing cerebellar nuclear neurons in vitro. Cerebellum 9, 67–76 (2010).

72. Gauck, V. & Jaeger, D. The control of rate and timing of spikes in the deep cerebellar nuclei by inhibition. J Neurosci 20, 3006–3016 (2000).

73. Hoebeek, F.E., Witter, L., Ruigrok, T.J. & De Zeeuw, C.I. Differential olivo-cerebellar cortical control of rebound activity in the cerebellar nuclei. Proc Natl Acad Sci U S A 107, 8410–8415 (2010).

74. Person, A.L. & Raman, I.M. Synchrony and neural coding in cerebellar circuits. Front Neural Circuits 6, 97 (2012).

75. Welsh, J.P., Lang, E.J., Suglhara, I. & Llinas, R. Dynamic organization of motor control within the olivocerebellar system. Nature 374, 453–457 (1995).

76. Boehme, R., Uebele, V.N., Renger, J.J. & Pedroarena, C. Rebound excitation triggered by synaptic inhibition in cerebellar nuclear neurons is suppressed by selective T-type calcium channel block. J Neurophysiol 106, 2653–2661 (2011).

77. Salinas, E. & Thier, P. Gain modulation: a major computational principle of the central nervous system. Neuron 27, 15–21 (2000).

78. Salinas, E. Fast remapping of sensory stimuli onto motor actions on the basis of contextual modulation. J Neurosci 24, 1113–1118 (2004).

79. Titley, H.K., Brunel, N. & Hansel, C. Toward a Neurocentric View of Learning. Neuron 95, 19–32 (2017).

